# Foldcomp: a library and format for compressing and indexing large protein structure sets

**DOI:** 10.1101/2022.12.09.519715

**Authors:** Hyunbin Kim, Milot Mirdita, Martin Steinegger

## Abstract

Highly accurate protein structure predictors have generated hundreds of millions of protein structures; these pose a challenge in terms of storage and processing. Here we present Foldcomp, a novel lossy structure compression algorithm and indexing system to address this challenge. By using a combination of internal and cartesian coordinates and a bi-directional NeRF-based strategy, Foldcomp improves the compression ratio by a factor of 3 compared to the next best method. Its reconstruction error of 0.08Å is comparable to the best lossy compressor. It is 5 times faster than the next fastest compressor and competes with the fastest decompressors. With its multi-threading implementation and a Python interface that allows for easy database downloads and efficient querying of protein structures by accession, Foldcomp is a powerful tool for managing and analyzing large collections of protein structures.

**Availability:** Foldcomp is a free open-source library and command-line software available for Linux, macOS and Windows at https://foldcomp.foldseek.com. Foldcomp provides the AlphaFold Swiss-Prot (2.9GB), TrEMBL (1.1TB) and ESMatlas HQ (114GB) database ready-for-download.

## INTRODUCTION

Fast and highly accurate structure prediction methods, such as AlphaFold2 [1] and ESMFold [2], have generated an avalanche of publicly available protein structures. The AlphaFold database [3] and ESMatlas [4] contain over 214 million and 617 million predicted structures in PDB format, respectively. A compressed local copy would require 25 and 15 TB storage, respectively. These databases are biological treasure troves but analyzing them is challenging due to these technical aspects.

The PDB or mmCIF [5] formats store protein structures as atom records in an 80-byte columnar plain-text format that includes the cartesian coordinates. Various strategies [6] have been proposed to deal with the growth of protein structure databases, including general-purpose compressors like Gzip and data-record-specific encodings like BinaryCIF [7] and MMTF [8]. PIC [9] transforms 3D coordinates into a lossy 2D image-like format and applies the PNG-image compression algorithm. Specialized formats for molecular trajectories [10] have also been developed to compress different states of a same molecule.

Here we present Foldcomp, a software and library that implements a novel algorithm to compress PDB/mmCIF using anchored internal coordinates combined with an indexing strategy to store large structural sets. We provide a command-line interface, a library/API for inclusion in other projects, and a Python interface to (de)compress and efficiently load user-selected entries in sequential- or random-access order.

## METHODS

Foldcomp’s workflow and file format is illustrated in **Fig. 1 a,b**. *Input* Foldcomp compresses PDB/mmCIF files stored in various input formats, such as individual-, directories of-, or optionally compressed tar-archives of PDB/mmCIF files. It returns compressed binary files (fcz format) in an individual directory, tar-archive, or Foldcomp database.

**FIG. 1:**
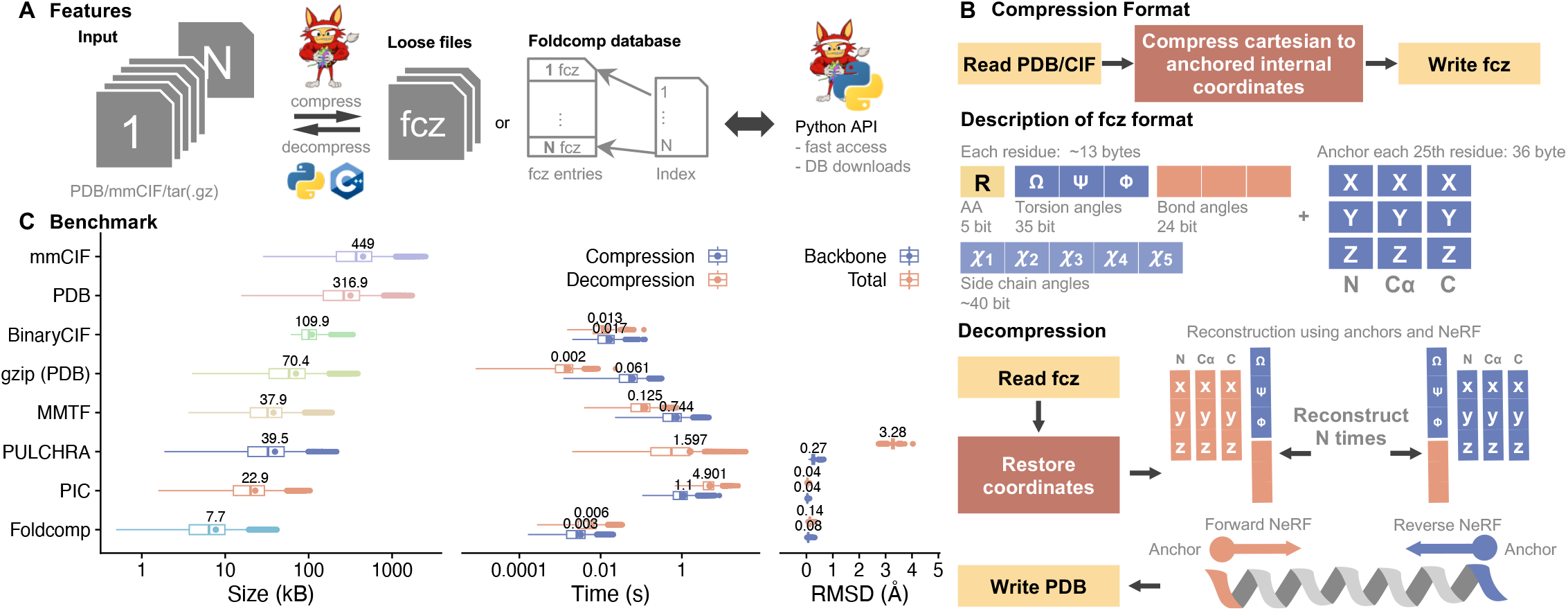
(**A**) Foldcomp is a library to compress, store and index protein structures. Foldcomp is written in C++ and comes with a command line and Python interface to compress, decompress, and access structures. (**B**) *Compression* takes 3D atom coordinates stored in PDB/mmCIF format as input and calculates and stores all internal coordinates, backbone torsions, and bond angles, and additionally, for every Nth residue (by default every 25th) the 3D atom coordinates for the N, C, and C-alpha atoms as anchor coordinates in its fcz format. By using anchors, we can prevent the accumulation of decoding errors. *Decompression* uses the anchor coordinates and internal coordinates to reconstruct the 3D atom coordinates by first extending the coordinates from N-terminal to C-terminal (forward) and then from C-terminal to N-terminal (backward) using Natural Extension Reference Frames (NeRF; [14]), followed by averaging the coordinates in between. Averaging reduces the reconstruction error by approximately a factor of two. (**C**) Comparison of file size (left), compression/decompression speed (middle), and backbone/all-atoms reconstruction error (right) for lossless and lossy protein structure compressors using the *Saccharomyces cerevisiae* proteome from the AlphaFold DB.

### Index

All compressed entries are concatenated and stored in a single file. We keep track of the entry identifier, start position, and length in separate plain text files. This format is compatible with the MMseqs2 [11] database format, which was initially inspired by the ffindex database format (unpublished). We added support for Foldcomp databases in Foldseek [12].

### Python

Foldcomp’s Python interface can be installed using pip install foldcomp. We provide functionality to download prebuilt databases, compress and decompress, and iterate.

## RESULTS

We compared Foldcomp to state-of-the-art software (**Fig. 1c**) using the *Saccharomyces cerevisiae* proteome from the AlphaFold DB v4. Among the compressors tested (PIC, PULCHRA [13], MMTF-python, Ciftools-java, and Gzip; see **Benchmark**), Fold-comp was the most efficient in terms of speed and size. It required 0.006 and 0.003 seconds for compression and decompression, respectively, and had a size of 7.7kb while maintaining one of the lowest reconstruction errors of 0.08Å and 0.14Å for backbone and all-atoms among the lossy compressors. The measurements were repeated and averaged over five trials. Using 16 threads reduced the time for compression and decompression to 1.617 and 2.532 seconds, respectively, resulting in a speed-up of 13x compared to single-thread.

### Databases

We provide a Foldcomp version of the AlphaFold database (v4) Swiss-Prot, TrEMBL and ESMatlas high-quality requiring 2.9GB, 1.1TB and 114GB, respectively. This is a magnitude smaller than the original size. Our databases are hosted on Cloud-Flare R2 for fast downloads and can be easily accessed through the Python interface.

## LIMITATIONS

Currently, Foldcomp only supports protein structures without missing residues. We plan to extend the format to deal with discontinuities in the future. Foldcomp is not meant to replace the PDB/mmCIF format, since these contain valuable metainformation that is discarded by Foldcomp.

## CONCLUSION

Foldcomp’s high speed combined with its novel algorithm to efficiently compress, and index structures will enable researchers to easily explore large collections of predicted protein structures on consumer hardware. We anticipate that easy access to billions of predicted protein structures will advance the field of protein structure analysis.

## ACKNOWLEDGEMENT

We thank Johannes Söding for the discussions and Do-Yoon Kim for creating the Foldseek logo. The work was supported by the National Research Foundation of Korea, grants [2019R1A6A1-A10073437, 2020M3A9G7103933, 2021R1C1C102065, 2021M3A9-I4021220], Samsung DS research fund and the Creative-Pioneering Researchers Program through Seoul National University.

## BENCHMARK

To benchmark protein structure compression tools, we used predicted structures of *Saccharomyces cerevisiae* proteome from AlphaFold database version 4 available at https://ftp.ebi.ac.uk/pub/databases/alphafold/latest/UP000002311_559292_YEAST_v4.tar. **Table I** shows the databases we used in this paper. All runtimes were measured on a server with an AMD EPYC 7702P 64-core CPU and 1TB RAM. All compression/decompression measurements are available in **Supplementary Data 1**.

**TABLE I:** Structure databases used in this manuscript.

| Database | Subset | Version | Entries | Size |
| --- | --- | --- | --- | --- |
| AlphaFoldDB | YEAST | v4 | 6039 | 1.8 GB |
| AlphaFoldDB | TrEMBL | v4 | 214 M | 23 TB |
| AlphaFoldDB | SwissProt | v3 | 542378 | 26 GB |
| ESMatlas | HQ | v0 | 37 M | 4.3 TB |

All benchmarks described were conducted on a RAM disk (/dev/shm) to minimize the effect of I/O operations for all the tools. **Table II and III** show runtime measurements executed on an NVMe SSD and HDD. These show similar runtimes to the in-RAM runtimes except for higher sys time.

**TABLE II:**
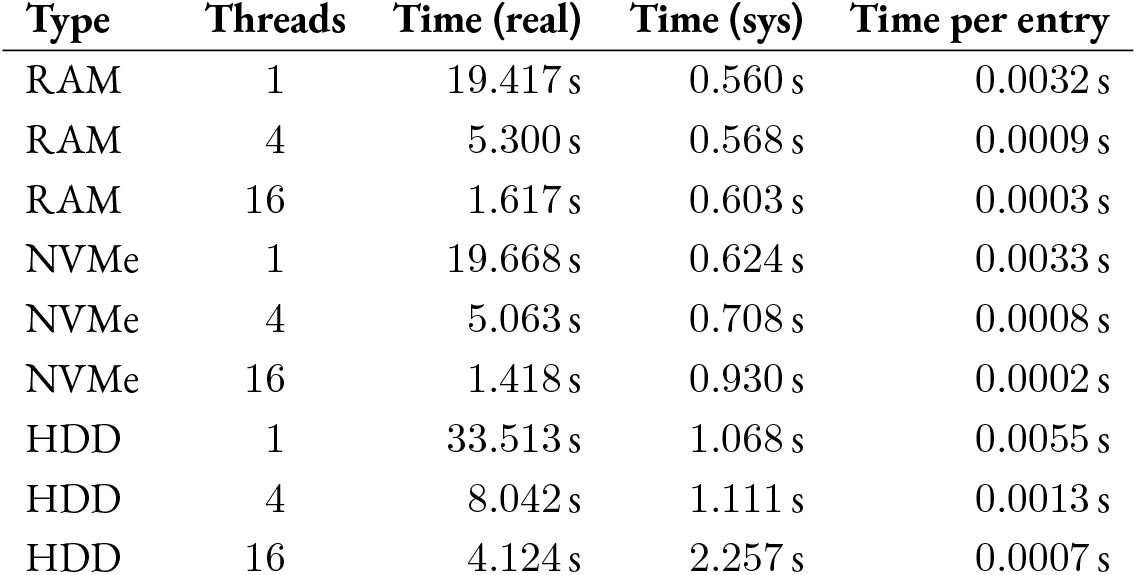
Benchmark of Foldcomp on compressing AlphaFoldDB-YEAST dataset with different thread counts and on different disk types.

**TABLE III:**
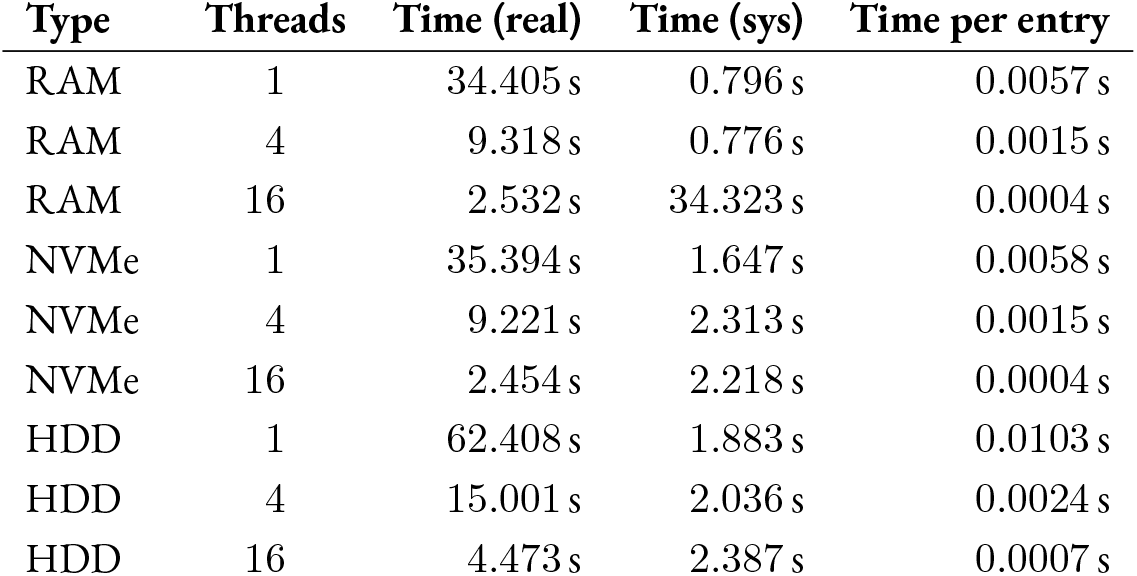
Benchmark of Foldcomp on decompressing AlphaFoldDB-YEAST dataset with different thread counts and on different disk types.

As Foldcomp supports batch mode for compression and decompression, we also used batch mode for the other compressors, if possible. For the tools that do not have a batch mode, we either implemented a batch mode or used GNU parallel [1].

*Foldcomp* was run in batch mode, which is automatically activated if a directory with PDB/mmCIF files is given as an input (or activated if a tar file containing PDB/mmCIF files is given). For time measurements, we use the `--time flag`, which measures the runtime of functions for compression and decompression with `std::chrono::high_resolution_clock`. **Fig. 2** shows the longest protein in AlphaFold-YEAST decompressed by Foldcomp, and superposed to the original PDB file using ChimeraX [2].

**FIG. 2:**
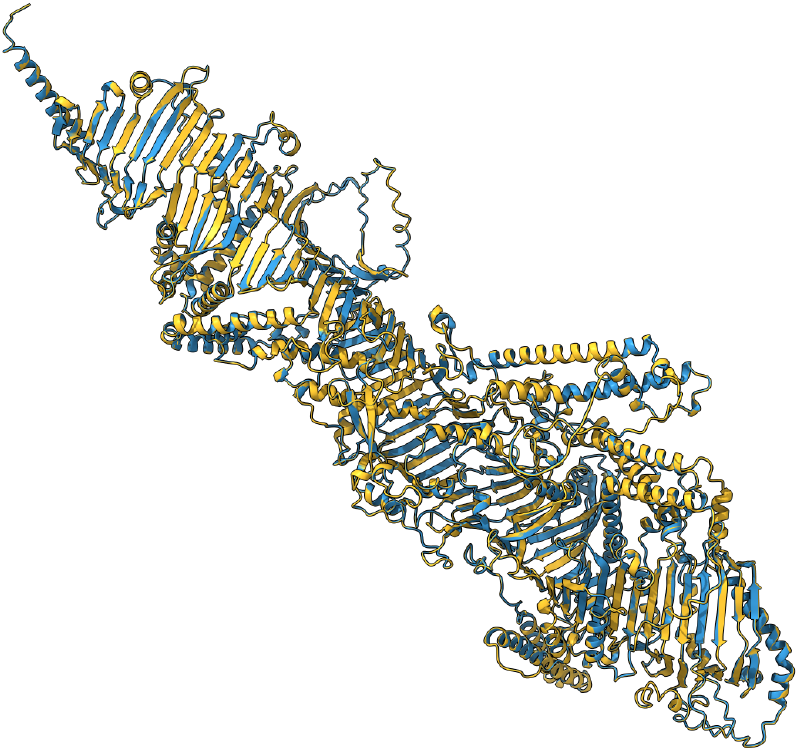
FMP27 (Q06179) is a mitochondrial protein with 2,628 aminoacids and 21,382 atoms, and also the largest protein in the AlphaFoldDB Yeast dataset. Saving this structure in `fcz` format requires 41.6 KB instead of 1.75 MB (PDB) with backbone RMSD of 0.079Å, and all-atoms RMSD of 0.14Å.

*Gzip* was executed using the recursive (`-r`) option for batch running and we added timestamps using `ts` from `moreutils` [3] to each entry to measure runtimes.

*PIC* was executed using the recursive (`-r`) option. To cut the total runtime, we split the input directory into the number of processes to run and applied `parallel`. We applied the `-rk` flag for compression and -rdk flag for decompression. During the batch execution loop, we measured the runtimes of `PICCompress` and `PICDecompress` with `time.process_time`.

*CIFTtools* is written in Java and has APIs to read/write mmCIF and binary CIF formats. As it lacks a command line interface for batch execution, we wrote `CompressBatch.java` and `DecompressBatch.java` to benchmark. We measured the runtime of `CifIO.readFromPath`, and either `CifIO.writeBinary` (compression) or `CifIO.writeText` (decompression) with `System.nanoTime`.

*MMTF* has been integrated into BioPython’s [4] `Bio.PDB API`. Using the API, we wrote `mmtf-batch.py` to benchmark. In the compression step, mmCIF files were loaded with `MMCIFParser` and MMTF files were written with `MMTFParser`. `MMTFIO` and `PDBIO` were used to read MMTF files and write PDB output during decompression. We measured runtime with `time.process_time`.

*PULCHRA* reconstructs whole peptides from C-alpha atoms using an optimization procedure and was not originally developed as a compression tool. We can achieve size reduction of PDB files by discarding all non-C-alpha atoms. As this is not an actual compression procedure, we have omitted the compression time number. The compression runtime number in the raw data table is based on grep-ping C-alpha atoms in PDB files. We used the runtimes reported by Pulchra.

Scripts used to benchmark are available at: https://github.com/steineggerlab/foldcomp-analysis. Used software versions for all tools are listed in **Table 4**, included libraries used to generate **Fig. 1c**.

**TABLE IV:**
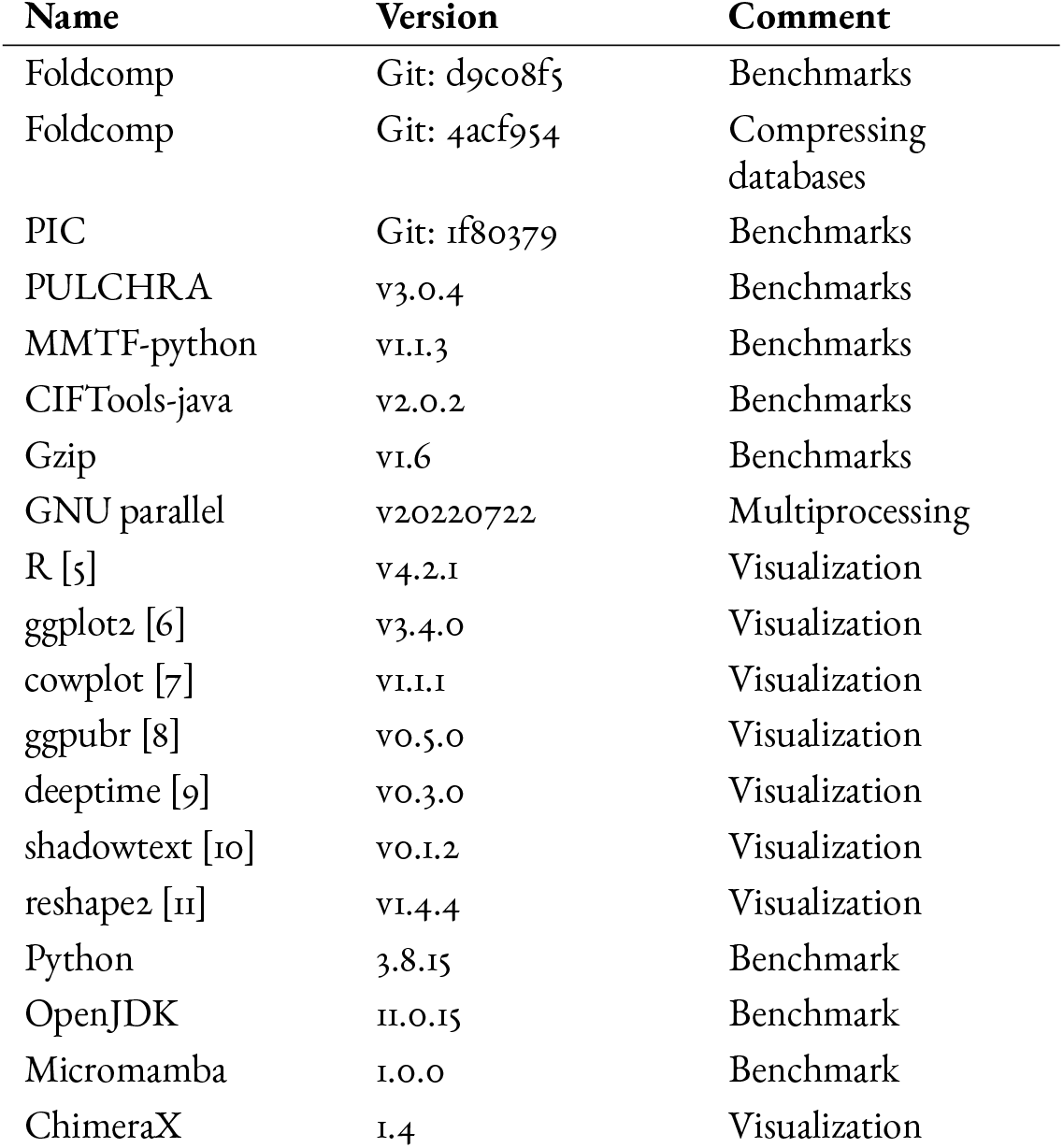
Software versions used in this manuscript.

